# Loss of the lncRNA *SOX1-OT* promotes p53-dependent cell-cycle arrest in astrocytes

**DOI:** 10.64898/2026.07.03.736275

**Authors:** Ulrike Fuchs, Sophie Schröder, Tonatiuh Pena, Dennis M Krüger, Susanne Burkhardt, Anna-Lena Schütz, Nina Hempel, Verena Gisa, Tarannom Taghavi, Jonathan Cortes, Michael Gertig, Ivana Delalle, Farahnaz Sananbenesi, André Fischer

## Abstract

Long non-coding RNAs (lncRNAs) are increasingly recognized as regulators of brain cell function, but their roles in astrocyte biology and neurodegeneration remain poorly understood. Here, we identify *Sox1ot/SOX1-OT* as a conserved, brain-enriched lncRNA that is downregulated in Alzheimer’s disease and in reactive astrocyte states. Antisense oligonucleotide-mediated depletion of *Sox1ot* in astrocytes revealed a transcriptional program marked by activation of p53 target genes selectively associated with cell-cycle inhibitory pathways. Consistent with this, *Sox1ot* depletion enhanced p53 occupancy at target promoters such as *Cdkn1a*, increased *Cdkn1a* expression and levels of its protein product p21, and thereby induced G1 arrest and reduced astrocyte proliferation. In contrast, other canonical p53 outputs, including apoptosis and senescence, were not affected, indicating that *Sox1ot* selectively modulates distinct branches of p53 signaling. Notably, loss of *Sox1ot/SOX1-OT* was accompanied by impaired glutamate uptake, reduced lactate secretion, and altered astrocyte support functions, suggesting that these deficits arise as downstream consequences of the p53-dependent transcriptional shift rather than direct primary effects of *Sox1ot* loss. Together, these findings identify *SOX1-OT* as an astrocyte-enriched regulatory layer that constrains a p53-dependent cell-cycle program and highlight its role in shaping astrocyte state transitions in Alzheimer’s disease.

## Introduction

Astrocytes are essential regulators of brain homeostasis and neural circuit function and constitute the most abundant glial cell population in the brain ^1 2^. In response to injury, inflammation, and neurodegenerative diseases such as Alzheimer’s disease (AD), astrocytes undergo marked state changes. In these contexts, astrocytes adopt reactive states associated with impaired homeostatic functions, altered inflammatory signaling, and changes in cell-cycle-related pathways, reflecting their capacity to proliferate and remodel their transcriptional state in response to stress or inflammatory cues ^3 4 5^. However, the molecular mechanisms that regulate these astrocyte state transitions remain incompletely understood. Because these transitions can profoundly influence neuronal function, identifying the molecular regulators that control astrocyte state is an important step toward understanding mechanisms of brain disease.

Long non-coding RNAs (lncRNAs) are attractive candidates for such regulation because many show strong tissue and cell type specificity and can shape gene expression programs in a context-dependent manner ^6^. Although lncRNAs are increasingly recognized as important regulators of brain function ^7 8 9^, their roles in glial biology remain insufficiently characterized. This is particularly relevant for pathways governing astrocyte functions which become altered in disease-associated reactive states ^3^.

One pathway of interest in this context is Tumor Protein p53 (p53) signaling. p53 is a transcription factor that induces programs of cell-cycle arrest, senescence, and apoptosis through a network of p53-responsive genes and proteins that interact with numerous other cellular signaling pathways ^10^ ^11 12^. p53 has also been implicated in neurodegenerative diseases, including AD ^13^ and astrocytic p53 is increasingly recognized as a key driver of neurodegenerative disease pathogenesis ^14^. However, the regulatory network governing p53-mediated astrocyte states is still poorly characterized.

Here, we prioritized the lncRNA *SOX1-OT* for functional analysis because it is brain-enriched, shows conserved expression in astrocytes across humans and mice, and is downregulated in AD and in reactive astrocyte-like conditions. We investigated its function in mouse and human astrocytes and found that *Sox1ot* regulates astrocyte support functions and p53-dependent cell-cycle control without affecting cellular senescence or apoptosis. Together, these findings identify *SOX1-OT* as an lncRNA-based regulatory layer that may shape astrocyte-specific p53 activity and point to this pathway as a potential target for modulating astrocyte dysfunction in AD.

## Materials & Methods

### Animals

Animal experiments were approved by the local Animal Welfare Office of Goettingen University and the Lower Saxony State Office for Consumer Protection and Food Safety (AZ:AZ 22.00203). CD-1 mice were obtained from Janvier Labs (Janvier Labs, France), transgenic mice were from the APP/PS1 mouse strain B6-Tg(Thy1-APPswe; Thy1-PS1 L166P) ^15 16^. Mice were housed in standard cages with a 12-h dark and light cycle. Water and food were provided ad libitum.

### Human tissues

For the analysis of *SOX1OT* expression by qPCR in human brains, tissue samples (prefrontal cortex, BA9) from control (n = 4 females & 3 males; age = 79 ± 10 years) and AD patients (n = 4 females & 3 males; age = 81 ± 6 years, Braak & Braak stage IV) were obtained with ethical approval from the ethics committee of the University Medical Center Göttingen and upon informed consent from the Harvard Brain Tissue Resource Center (Boston, USA). Log2 fold change values for brain-enriched lncRNAs were obtained from the AGORA database, a multi-omic resource that integrates transcriptomic, proteomic, and metabolomic data from human postmortem brain samples across AD and control cohorts. For lncRNAs reported as differentially expressed by AGORA (L2FC |0.263|), L2FC values were averaged across those regions to determine concordant deregulation, and then averaged to yield a single representative L2FC value.

### Primary astrocyte culture

Primary astrocyte cultures were generated from postnatal day 1-2 mouse cortices and hippocampi as previously described ^8^. Briefly, tissue was dissociated with trypsin-EDTA (ThermoFisher, MA, USA), and cells from 2-3 pups were plated on poly-D-lysine-coated flasks in serum-containing DMEM (ThermoFisher), then shaken 7 days later to remove non-astrocytic cells and re-plated in Neurobasal Plus medium supplemented with B-27 Plus, GlutaMAX, penicillin/streptomycin (all ThermoFisher), and HB-EGF (Merck, Germany) at 15 000 cells/cm². Cultures were maintained at 37°C, 5% CO₂ with half-medium changes twice per week.

### Primary cortical neuron culture

Primary neuronal cultures were generated as previously described ^8^. In short, pregnant CD-1 mice were sacrificed using pentobarbital, and embryonic day 17 brains were rapidly dissected in ice-cold PBS (ThermoFisher) with meninges removed. Cortices were dissociated using the Papain Dissociation System (Worthington, NJ, USA) following the manufacturer’s instructions. Cells were resuspended in Neurobasal Plus medium supplemented with B-27 Plus, GlutaMAX, penicillin/streptomycin (all ThermoFisher), and seeded at densities of 50 000 cells/cm². For co-cultures, astrocyte-containing inserts were added to cortical neurons at day in vitro (DIV)14, with experiments conducted on DIV17. Cultures were maintained at 37°C, 5% CO₂ with a third-medium change every 2 days.

### Human iPSC-derived astrocytes

Human iPSC-derived astrocytes (Ncyte Astrocytes) were obtained from Ncardia (Ncardia, Netherlands). Following the manufacturer’s instructions, cells were thawed and cultured for 1-2 weeks before performing experiments.

### Stimulation of astrocytes

To mimic the activation of astrocytes by microglia ^17^, astrocytes were treated with a cytokine mixture of Il-1α (3 ng ml^−1^, Merck, I3901), TNF (30 ng ml^−1^, Cell Signaling Technology, MA, USA, 8902SF) and C1q (400 ng ml^−1^, MyBioSource, CA, USA, MBS143105) for 24 hours. To mimic the activation of astrocytes during Alzheimer’s disease, Amyloid beta 1-42 (Cayman Chemicals, MI, USA) was prepared as previously described ^8^ and protofibrils were added to astrocytes at a concentration of 10 µM for 24 hours.

### Antisense LNA Gapmers

Antisense LNA Gapmers targeting *Sox1ot* and *SOX1-OT* and negative controls (NC) were designed and purchased from Qiagen (Qiagen, Netherlands) having the following sequences:

NC: GCTCCCTTCAATCCAA

*Sox1ot (1)*: CACGCTAAGTTGAGAT

*Sox1ot (2)*: AGTGCTGACTAAAAGT

*SOX1-OT*: AAGTTTAGTCCTCGGCTCTG

Antisense LNA GapmeRs were designed using Qiagen’s GeneGlobe Antisense LNA GapmeR Custom Builder, which selects RNase H-dependent ASOs with no predicted RISC-associated off-target activity. The design tool evaluates potential off-targets by sequence alignment against spliced and unspliced ENSEMBL transcriptomes. To independently validate the specificity of the GapmeRs, we performed a BLAST alignment of the antisense sequences against the full mouse transcriptome to identify potential reverse-complement matches.

Transfection was performed using Lipofectamine RNAiMAX (ThermoFisher) according to the manufacturer’s instructions. Cells were treated on DIV12 and all experiments were performed on DIV14. Cell viability was measured using PrestoBlue (ThermoFisher).

### *Sox1ot* overexpression using CRISPRa

Plasmids containing the guide RNA (gRNA), a dCas9-VP64 and a tGFP reporter gene were obtained from Origene (GA217630) (Origene, MD, USA). A scramble gRNA (scramble) with no target in the genome was used as a negative control (GE100077). To increase gene expression, an enhancer vector (GE100056) was used in all cases. The gRNA for targeting Sox1ot was designed by Origene and had the following sequence:

#### gRNA-3: AAGACTCGCAAGACAATTTC

Primary astrocytes were transfected as previously described ^8^ using the Neon Nxt electroporation device (ThermoFisher). In short, 100 000 cells were electroporated using 0.5 µg CRISPRa plasmid and 0.1625 µg enhancer plasmid (1300 V, 20 ms, 2 pulses). Cells were seeded at a density of 50 000 cells/cm2. After 72 hours, functional experiments and RNA isolation were performed.

### Glutamate uptake

On DIV14, primary astrocytes were incubated with 100 µM glutamate for 1 hour as previously described ^8^. Supernatant was collected and the amount of remaining glutamate was determined using the Glutamate-Glo™ Assay (Promega, WI, USA) following the manufacturer’s instructions. Luminescence was recorded with a FLUOstar® Omega (BMG Labtech, Germany).

### Lactate secretion

To estimate lactate secretion, the complete medium was changed 24 hours before the endpoint of an experiment. The medium was collected and lactate concentrations were measured using the Lactate-Glo™ Assay (Promega) following the manufacturer’s instructions.

### Senescence

Cells were seeded in white 96-well plates and senescence-associated β-galactosidase (SA-β-gal) activity was quantified using the Beta-Glo Assay (Promega) according to the manufacturer’s protocol. In short, after equilibrating plates to room temperature, an equal volume of Beta-Glo reagent to culture medium was added, plates were mixed for 30 sec, and incubated for 30 min at room temperature in the dark before luminescence measurement on a FLUOStar Omega plate reader (BMG Labtech).

### Apoptosis

Apoptosis was quantified using the RealTime-Glo Annexin V Apoptosis and Necrosis Assay (Promega) following the manufacturer’s instructions. Cells were seeded in white 96-well plates and subjected to *Sox1ot* antisense LNA Gapmer knockdown. Then, prewarmed medium containing a 1× mixture of detection reagents was added. Fluorescence and luminescence were then recorded at 37°C at serial time points on a FLUOStar Omega plate reader (BMG Labtech) for up to 72 h, and background-subtracted signals were analyzed using GraphPad Prism.

### Intracellular calcium flux

Intracellular calcium responses were measured using the Fluo-4 Direct Calcium Assay Kit (ThermoFisher). Culture medium was reduced to 50 µl per well in 96-well plates, Fluo-4 loading solution containing probenecid was added, and cells were incubated for 30 min at 37°C before baseline fluorescence (RFU_Min_)was recorded at 494/516 nm on a FLUOStar Omega plate reader (BMG Labtech). Subsequently, L-glutamate (Abcam, MA, USA) was added to a final concentration of 100 µM, wells were rapidly mixed, and fluorescence was recorded again (RFU_Max_). Finally, the blank-corrected difference in relative fluorescence units (ΔRFU) was calculated using the following formula: ΔRFU = (RFU_Max_ − RFU_Min_)/RFU_Min_.

### Ki67 immunofluorescence proliferation assay

Cell proliferation was assessed by Ki67 immunofluorescence staining. Cells were seeded on 0.1% PEI-coated 8-chamber slides at 7 000 cells per chamber, fixed in 4% paraformaldehyde for 10 min at room temperature, and washed in PBS (ThermoFisher). Cells were permeabilized with 0.1% Triton X-100 (Merck) in PBS for 15 min, washed, and blocked for 1 h in PBS containing 2% BSA (Cell Signaling Technologies), 20% normal goat serum (NGS) (Abcam), and 0.1% Tween-20 (Merck). Slides were incubated overnight at 4°C with anti-Ki67 (Abcam) primary antibody diluted in 2% BSA and 5% NGS in PBS containing 0,1 % Tween20 (PBS-T), washed, and then incubated for 1 h at room temperature with the appropriate fluorophore-conjugated secondary antibody in 2% BSA and 1,5% NGS in PBS-T, followed by extensive washing. Nuclei were counterstained with DAPI (Merck) for 5 min, rinsed in PBS, mounted with ProLong Gold antifade mountant (LifeTechnologies, CA, USA). Images were acquired on a Keyence BZ-X800 microscope (Keyence, Japan) using 20x or 40x objectives.

### Multielectrode array

Multielectrode array (MEA) recordings were performed in the Maestro system (Axion), as previously described ^18^. *Sox1ot* knockdown primary astrocytes were seeded on top of primary cortical neurons at DIV10, and recordings took place every 24 hours for up to 3 days. A neuronal activity score (NAS) was calculated according to a recent publication ^19^.

### Cytoplasmic and nuclear fractionation

Cytoplasmic and nuclear fractions were prepared as previously described ^8^. In brief, DIV14 primary astrocytes were fractionated using the EZ prep lysis kit (Merck) according to the manufacturer’s protocol, supplemented with RNase inhibitor (Promega), followed by TRIzol LS (ThermoFisher) extraction and storage at -20°C.

### *In situ* hybridization combined with immunofluorescence

RNAscope Fluorescent Multiplex assays (Advanced Cell Diagnostics, CA, USA) combined with GFAP immunofluorescence was performed on 18 µm fresh frozen tissue sections per manufacturer’s instructions. Sections were fixed in 10% NBF, treated with H₂O₂, incubated with anti-GFAP (1:250, Abcam) or anti-NeuN (1:200, Abcam) and hybridized with Sox1ot probes. Then, probes were detected with TSA Plus Cy5 (1:750), counterstained with Alexa Fluor 555 anti-rabbit (1:1000, ThermoFisher) and DAPI, and mounted in ProLong Gold antifade mountant. Images were acquired on a Keyence BZ-X800 microscope using a 63x oil objective.

### RNA extraction

Total RNA was extracted from cells using TRI Reagent (Merck) and the RNA Clean & Concentrator-5 kit (Zymo Research, CA, USA) per manufacturer’s instructions. Concentrations were quantified by NanoDrop or Qubit (RNA HS Assay; ThermoFisher), and quality assessed by Bioanalyzer (Agilent, CA, USA) for sequencing samples.

### Library preparation and total RNA sequencing

300 ng total RNA using the SMARTer Stranded Total RNA Sample Prep Kit-HI Mammalian (Takara Bio, Japan), amplified for 13 cycles, and sequenced on a NextSeq 2000 (Illumina, CA, USA; 50 bp single-end).

### Bioinformatic analysis

Raw reads were processed with bcl2fastq (v2.20.2), quality-checked with FastQC (v0.11.5), aligned to mm10 using STAR (v2.5.2b), and quantified with featureCounts (v1.5.1). Differential expression analysis was performed using DESeq2 (v1.38.3) ^20^ with RUVSeq (v1.32.0) correction ^21^, followed by GO analysis with clusterProfiler (v4.6.0) ^22^ and transcription factor enrichment via Enrichr (https://maayanlab.cloud/Enrichr/).

### cDNA, RT-qPCR

cDNA was synthesized from 200-400 ng RNA using the Transcriptor First Strand Kit (Roche, Switzerland) with random hexamers. 2 ng of cDNA per sample were used for RT-qPCR with LightCycler 480 SYBR Master Mix (Roche) in duplicates on a LightCycler 480 (Roche). Expression of non-coding and protein-coding genes was normalized to 18S using the 2−ΔΔCt method ^23^. Primer sequences are listed in **Supplemental Table 11.**

### Western blot

Cytoplasmic and nuclear fractions of primary astrocytes were prepared and lysed in RIPA buffer (ThermoFisher) with protease inhibitors, quantified by BCA assay (ThermoFisher). After 10 min of heat denaturation in Laemmli buffer containing beta mercaptoethanol at 95°C, 10 µg of protein was separated on 4-20% TGX gels (Bio-Rad). Proteins were transferred to low-fluorescence PVDF using Trans-Blot Turbo (Bio-Rad Laboratories, CA, USA), blocked in 5% BSA/PBS-T, and probed overnight with anti-p21 (1:1000; ThermoFisher), anti-p53 (1:500; Cell Signaling Technologies), or anti-Lamin B1 (1:1000; Proteintech, IL, USA) (antibodies are listed in **Supplemental Table 12**). Fluorescent IRDye secondaries (LI-COR Biosciences, NE, USA) enabled detection on Odyssey DLx (LI-COR). Quantification was done in ImageJ.

### Chromatin immunoprecipitation with reference exogenous chromatin (ChIP-Rx)

ChIP-Rx was performed using the ChIP-Rx kit (Active Motif, CA, USA) per manufacturer’s instructions, to normalize for varying protein input amounts across astrocyte samples. DIV14 astrocytes treated with control or Sox1ot ASOs were cross-linked using 1% paraformaldehyde (Merck), quenched by glycine (Roth, Germany), and chromatin sheared by sonication at 4°C with a Bioruptor Plus (Diagenode, NJ, USA) in RIPA buffer containing 10% SDS. Immunoprecipitations used 10 µg chromatin and 2 µg Drosophila Spike-in chromatin (Active Motif) with 4 µg anti-p53 (Cell Signaling Technologies) or IgG (Cell Signaling Technologies) with 8 ng Drosophila Spike-in antibody (Active Motif), immune complexes were captured with protein A/G magnetic beads (ThermoFisher) as previously described ^8^; washed sequentially, and DNA for qPCR analysis was purified using the ZYMO ChIP Clean and Concentrator Kit (Zymo Research).

### Cell cycle analysis

Cell cycle distribution was analyzed by propidium iodide (PI) (ThermoFisher) staining and flow cytometry. Cells were synchronized by 48 h serum deprivation to arrest them in G1, followed by 24 h of serum re-addition and 24 h knockdown where indicated. Cells were detached with 0.25% trypsin/EDTA (ThermoFisher), pelleted, and fixed by resuspension in ice-cold 70% ethanol, followed by 60 min incubation on ice with intermittent vortexing. After washing with PBS, cells were resuspended in a PI staining solution containing 50 µg/ml PI, 0.1% Triton X-100, and incubated for 30 min at 37°C in the dark with gentle agitation. Finally, cells were washed, resuspended in ice-cold PBS, and analyzed on a BD FACS Diva (BD Biosciences, CA, USA) equipped with an 85-µm nozzle using forward and side scatter to gate singlets and PerCP-Cy5-5 (PI) signal, followed by gating to determine the proportion of cells in G0/G1, S, and G2/M phases.

### Statistical analysis

Statistical analysis was performed in GraphPad Prism 9, with data presented as mean +SEM unless otherwise indicated. Two-tailed unpaired t-tests or one-way ANOVA with Tukey’s post-hoc were used as appropriate. GO and pathway enrichments employed Fisher’s exact test with Benjamini-Hochberg correction.

## Results

### *SOX1-OT* is a brain-enriched lncRNA downregulated in AD

To identify lncRNAs enriched in the human brain and dysregulated in neurodegenerative disorders, namely in AD could represent a promising approach to find novel drug targets. Moreover, RNA therapeutics is an emerging area in neurodegenerative diseases with great potential to expand the scope of drug discovery, promising rapid translation into first in human clinical trials ^24^. However, still little is known about the role of lncRNAs in the adult brain. Therefore, identifying candidate lncRNAs and beginning to elucidate their function was a key objective of the current study. To further elucidate the role of lncRNAs in the human brain and AD, we first employed a dataset based on the RNA Atlas ^25 8 9^ to search for brain-enriched lncRNAs. We identified 32 candidates that exhibited more than 20-fold enriched expression in the brain compared to other human tissues **(Fig. 1A Supplemental Table 1)**. Next, we asked how many of these lncRNAs were significantly deregulated in postmortem brain tissue from AD patients using the AGORA database (https://agora.adknowledgeportal.org/), which includes over 1000 postmortem brain samples from control individuals and AD patients. We defined AD-associated candidates as lncRNAs showing at least a 20% concordant expression change across the measured brain regions, corresponding to an absolute log2 fold change of 0.26. This analysis revealed 15 candidates, of which three were evolutionarily conserved and had confirmed rodent homologues, namely *DLX6-AS1/Dlx6os1*, *SOX2-OT/Sox2ot*, and *SOX1-OT/Sox1ot* **(Supplemental Table 2)**. We focused on evolutionary conservation lncRNAs for further prioritization, since conservation suggests functional relevance.

**Figure 1.**
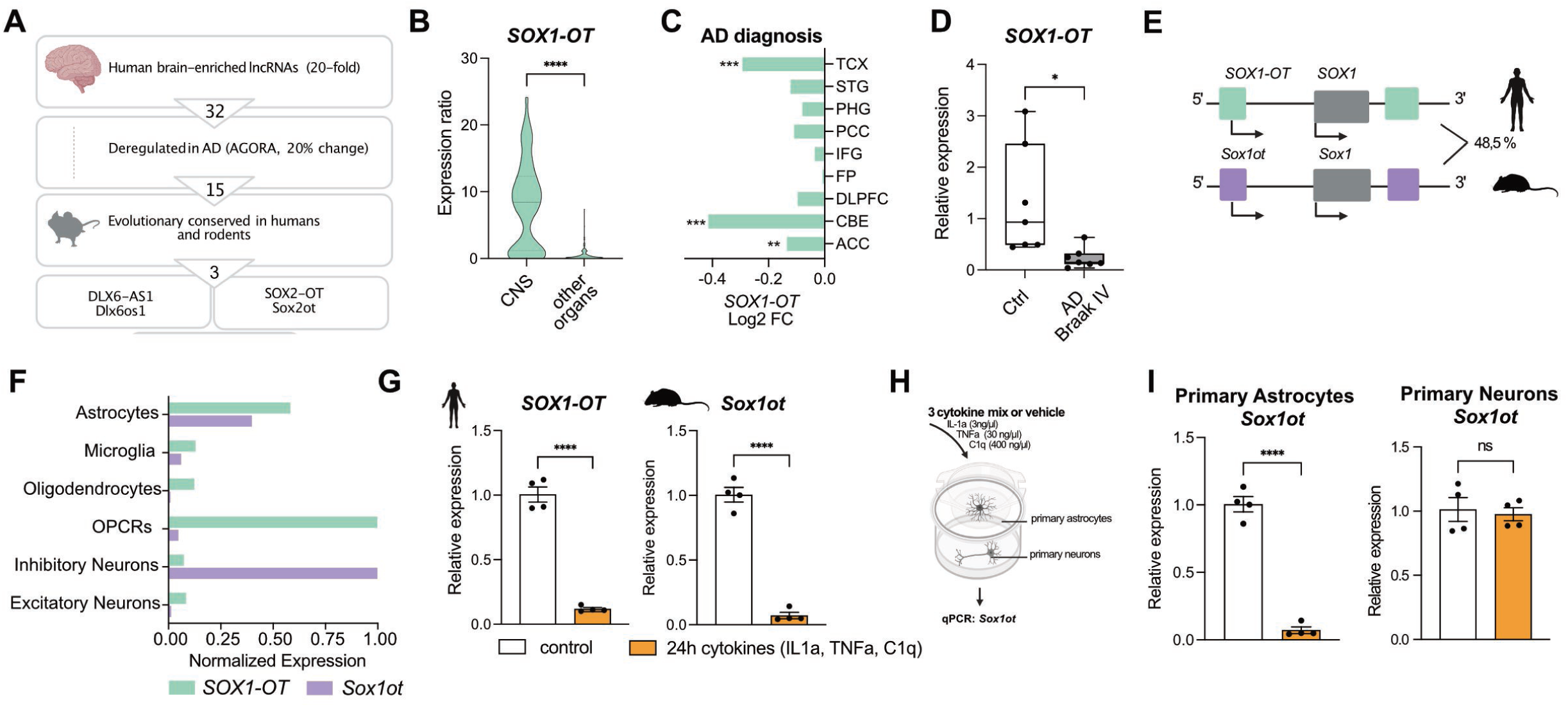
*SOX1-OT* is a brain-enriched lncRNA downregulated in Alzheimer’s disease. **A** Schematic overview of filtering criteria for lncRNA candidate selection, AD Alzheimer’s disease. **B** Expression of *SOX1-OT* in different human tissues (depicted is ratio of CNS compared to 24 other organs) obtained from ^25^. **C** Bar chart depicting Log2 Fold change (FC) of *SOX1-OT* in different brain regions in Alzheimer’s disease (AD) patients compared to controls based on data from the Agora database (https://agora.adknowledgeportal.org/), (\**P* < 0.05, \*\**P* < 0.01, \*\*\**P* < 0.001). **D** Bar plot showing qPCR results of *SOX1-OT* in human postmortem brain samples of Braak Stage IV compared to non-demented controls (unpaired *t-*Test, \**P* < 0.05). **E** Schematic representation of Sox1ot genomic locus in humans and rodents. Percentage indicating global sequence similarity. **F** Bar plot showing normalized expression of human *SOX1-OT* and mouse *Sox1ot* in different brain cell types, obtained from snucRNAseq data (iCELL8 technology) from healthy human brains (prefrontal cortex, BA9) ^8^ and snucRNAseq data (10X technology) from healthy mice (hippocampus) ^48^. **G** Bar plots showing qPCR result for human *SOX1-OT* or mouse *Sox1ot* after treating human induced pluripotent stem cell-derived astrocytes (left panel) or primary mouse astrocytes (right panel) with the cytokine mix IL1a, TNFa, C1q (cytokines) for 24 hours (unpaired *t-*Test; \*\*\*\**P* < 0.0001). **H** Schematic illustration of astrocyte - neuron co-culture system using 1 µM membrane inserts and subsequent cytokine treatment to induce inflammation. **I** Bar plots showing *Sox1ot* expression upon 24 hour treatment with the cytokines IL1a, TNFa, C1q within the insert co-culture system illustrated in (H) in primary astrocytes (left) or primary cortical neurons (right). Gene expression was normalised to 18S (unpaired *t* test, \*\*\*\**P* < 0.0001, ns = not significant). Error bars represent SEM.

Compared with the protein-coding genome, the function of most lncRNAs remains poorly understood. However, *DLX6-AS1* and *SOX2-OT* are already comparatively well-studied lncRNAs with links to neural development, cancer, and disease-associated processes, including emerging links to AD and related neurodegenerative conditions ^26 27 28^ **(Supplemental Table 2)**. In contrast, *SOX1-OT* remains largely uncharacterized. Only six studies have addressed its function to date **(Supplemental Table 2)**, linking it to neural differentiation ^29^, cancer-associated processes ^30 31^, and genetic association studies, including polymorphisms associated with cleft lip ^32 33^. Importantly, *SOX1-OT* has not been linked to adult brain function or neurodegenerative disease. We therefore selected *SOX1-OT/Sox1ot* as a conserved, AD-associated, and poorly characterized lncRNA candidate for functional analysis.

In support of this prioritization, *SOX1-OT* was strongly enriched in the CNS **(Fig. 1B)** and showed decreased expression in AD patient brains according to the AGORA database. Specifically, **SOX1-OT** expression was significantly reduced in the temporal cortex, cerebellum, and anterior cingulate cortex of AD patients **(Fig. 1C)**. We further validated this finding by qPCR analysis of postmortem prefrontal cortex samples (BA9) from healthy controls and patients with AD pathology at Braak stage IV **(Fig. 1D)**. Since *SOX1-OT* is conserved between humans and mice as measured by synteny **(Fig. 1E)**, we could investigate its function in murine and human model systems.

To further explore this, we first examined the cell type-specific expression of *SOX1-OT* in single-nucleus (sn) RNA-seq data from the prefrontal cortex (PFC) of neurologically healthy individuals and found that it is primarily expressed in astrocytes and oligodendrocyte precursor cells (OPCs) **(Fig. 1F, Fig. S1A)**. In the mouse brain, *Sox1ot* expression was highest in inhibitory neurons and astrocytes **(Fig. 1F; Fig. S1B)**. Although these data indicate that *SOX1-OT* is not confined to a single neural cell type, astrocytes were the only cell type to show consistently high expression of both *SOX1-OT* and *Sox1ot* across human and mouse brain samples **(Fig. 1F)**. This conserved cell type-specific expression pattern suggests that *SOX1-OT/Sox1ot* may have an important function in astrocytes.

To investigate whether *Sox1ot/SOX1-OT* is deregulated in astrocytes in the context of AD, we analyzed snRNA-seq data from the ROSMAP cohort ^34^. However, because these data were generated using a 3′-end poly(A)-selected 10x Genomics platform (10x Genomics, CA, USA), lncRNA expression was substantially underrepresented, and *SOX1-OT* could not be sufficiently detected **(Fig. S2)**. Please note that the dataset shown in Fig. 1F was generated using the iCELL8 platform (Takara, Japan) with a full-length total RNA-seq approach, which provides improved detection of lncRNAs ^8^. Since such data is not yet available from human AD brains, we decided to model astrocyte activation *in vitro* using a cytokine mix consisting of interleukin-1 alpha (IL-1α), tumor necrosis factor alpha (TNF-α), and complement component 1q (C1q), which induces reactive astrocyte gene expression changes resembling those observed in AD patients ^17^. Under these conditions, both mouse *Sox1ot* and human *SOX1-OT* levels were significantly reduced in primary mouse astrocytes and human induced pluripotent stem cell (iPSC)-derived astrocytes after 24 h of treatment with the cytokine mix **(Fig. 1G)**. When we repeated this experiment in a transwell culture system, in which primary neurons and astrocytes were cultured and exposed to the three-cytokine mixture. Of note, *Sox1ot* expression remained unchanged in primary neurons but was markedly reduced in astrocytes **(Fig. 1H, I)**. This was recapitulated, however, to a much lesser extent, when the same system was subjected to amyloid beta1-42 protofibrils **(Fig. S3A)**, suggesting that amyloid pathology can affect *Sox1ot* levels but is unlikely to be the sole or dominant direct driver of *Sox1ot* downregulation in astrocytes. Consistently, we observed a non-significant trend toward reduced *Sox1ot* expression in the dentate gyrus of 4-month-old APP/PS1 mice, representing an early stage of pathology **(Fig. S3B)**. Together, these findings support a model in which amyloid pathology may contribute to *Sox1ot* deregulation indirectly, potentially through microglial activation and subsequent inflammatory signaling to astrocytes.

This is in line with recent studies suggesting that amyloid beta acts synergistically with cytokines to promote pro-inflammatory astrocyte activation, rather than serving as the sole driver ^35^. Since the IL-1α/TNF-α/C1q cytokine mixture models a pathological microglial secretory state associated with AD ^17^, our findings suggest that AD-related inflammatory cues, such as reactive microgliosis, may contribute to reduced SOX1-OT/Sox1ot expression in astrocytes, but not in neurons. Taken together, these data further support the idea that astrocytic SOX1-OT/Sox1ot is deregulated during AD pathogenesis. We therefore focused our functional analyses on SOX1-OT/Sox1ot in astrocytes, while acknowledging that this lncRNA may also have important functions in other neural cell types, including OPCs and neurons.

### *Sox1ot* knockdown impairs astrocyte support functions for neurons

To characterize the cellular changes induced by the reduction of *Sox1ot* in astrocytes, as observed in AD, we performed Gapmer antisense oligonucleotide (Gapmer-ASO)-mediated knockdown (KD) of *Sox1ot* in primary astrocytes **(Fig. 2A)** and assessed key astrocyte functions 48 h after KD. Gapmer ASOs are single-stranded antisense oligonucleotides that mediate RNase H-dependent degradation of nuclear RNA transcripts. As a negative control, we used Gapmer ASOs with no known target in the genome.

**Figure 2.**
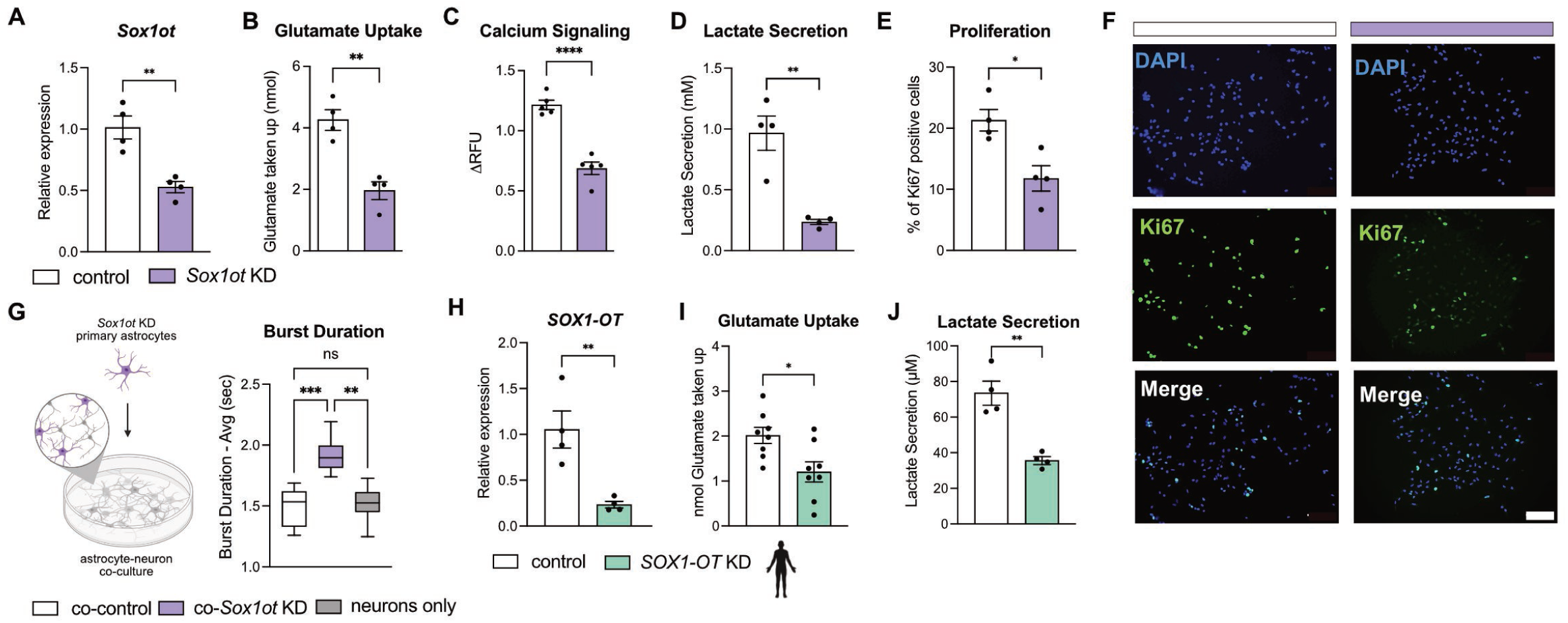
*Sox1ot* knockdown impairs astrocyte support functions for neurons in mouse and human. **A** Bar plot showing qPCR results for *Sox1ot* after treating primary astrocytes with Gapmers to knock down (KD) *Sox1ot*. RNA was collected 48 hours after addition of Sox1ot or control Gapmers (unpaired *t-*Test; \*\**P* < 0.01). Gene expression was normalized to 18S. **B** Bar blot showing results of the glutamate uptake assay in primary astrocytes 48 hours after *Sox1ot* knock down in comparison to control Gapmers (unpaired *t-*Test; \*\**P* < 0.01). **C** Bar blot showing results of the calcium signaling response to L-Glutamate (100 µM) in primary astrocytes 48 hours after *Sox1ot* knock down in comparison to control Gapmers (unpaired *t-*Test; \*\*\*\**P* < 0.0001). **D** Bar blot showing results of the lactate secretion assay in primary astrocytes 48 hours after *Sox1ot* knock down in comparison to control Gapmers (unpaired *t-*Test; \*\**P* < 0.01). **E** Bar chart showing the relative amount of proliferating primary astrocytes determined by Ki67 immunofluorescence imaging 48 hours after *Sox1ot* knock down in comparison to control Gapmers (unpaired *t-*Test; \**P* < 0.05). **F** Representative immunofluorescence images of (E) with anti-Ki67 antibody. Nuclei are stained with DAPI. Scale bar 100 µm. **G** Left panel: Schematic illustration of astrocyte-neuron co-culture. Knock down (KD) of *Sox1ot* was performed 24 hours before seeding of astrocytes on top of primary cortical neurons in a 1:5 astrocyte to neuron ratio. Right panel: Bar plot showing results of multielectrode assay recorded in primary cortical neurons cultured alone (neurons only) or co-cultured for 24 hours with *Sox1ot* depleted (co-Sox1ot KD) or control Gapmer-treated (co-control) primary astrocytes (one-way ANOVA; \*\**P* < 0.01, \*\*\**P* < 0.001, ns not significant). **H** Bar plot showing qPCR result for *SOX1-OT* after treating human induced pluripotent stem cell-derived astrocytes 48 hours with Gapmers to knock down (KD) human *SOX1-OT* (unpaired *t-*Test; \*\**P* < 0.01). Gene expression was normalised to 18S. **I** Bar blot showing results of the glutamate uptake assay in human induced pluripotent stem cell-derived astrocytes 48 hours after *SOX1-OT* knock down in comparison to control Gapmers (unpaired *t-*Test; \**P* < 0.05). **J** Bar blot showing results of the lactate secretion assay in human induced pluripotent stem cell-derived astrocytes 48 hours after *SOX1-OT* knock down in comparison to control Gapmers (unpaired *t-*Test; \*\**P* < 0.01). Error bars represent SEM.

*Sox1ot* KD led to significantly reduced glutamate uptake, a diminished intracellular calcium response to glutamate, decreased lactate secretion, and attenuated proliferation **(Fig. 2B–F)**. Next, we co-cultured cortical neurons with astrocytes subjected to *Sox1ot* KD or with corresponding control astrocytes and performed multielectrode array (MEA) recordings to assess neuronal network activity. We observed increased burst duration in neurons co-cultured with *Sox1ot*-KD astrocytes compared with control conditions **(Fig. 2G)**. This effect is consistent with the reduced glutamate uptake observed in *Sox1ot-*KD astrocytes. Furthermore, knockdown of human *SOX1-OT* impaired glutamate uptake and lactate secretion in iPSC-derived astrocytes, mirroring the effects observed in primary mouse astrocytes **(Fig. 2I,J).** Overall, these findings demonstrate that *SOX1-OT* plays a conserved functional role in astrocytes, and that its depletion impairs key astrocytic processes such as glutamate uptake and metabolic support.

To gain insight into its potential regulatory mechanisms, we examined the subcellular localization of *Sox1ot* in primary astrocytes using cellular fractionation and RNAscope imaging. Both methods revealed that *Sox1ot* is present in both the nucleus and the cytoplasm **(Fig. 3A)**, suggesting potential roles in the regulation of gene expression at both the transcriptional and post-transcriptional levels.

**Figure 3.**
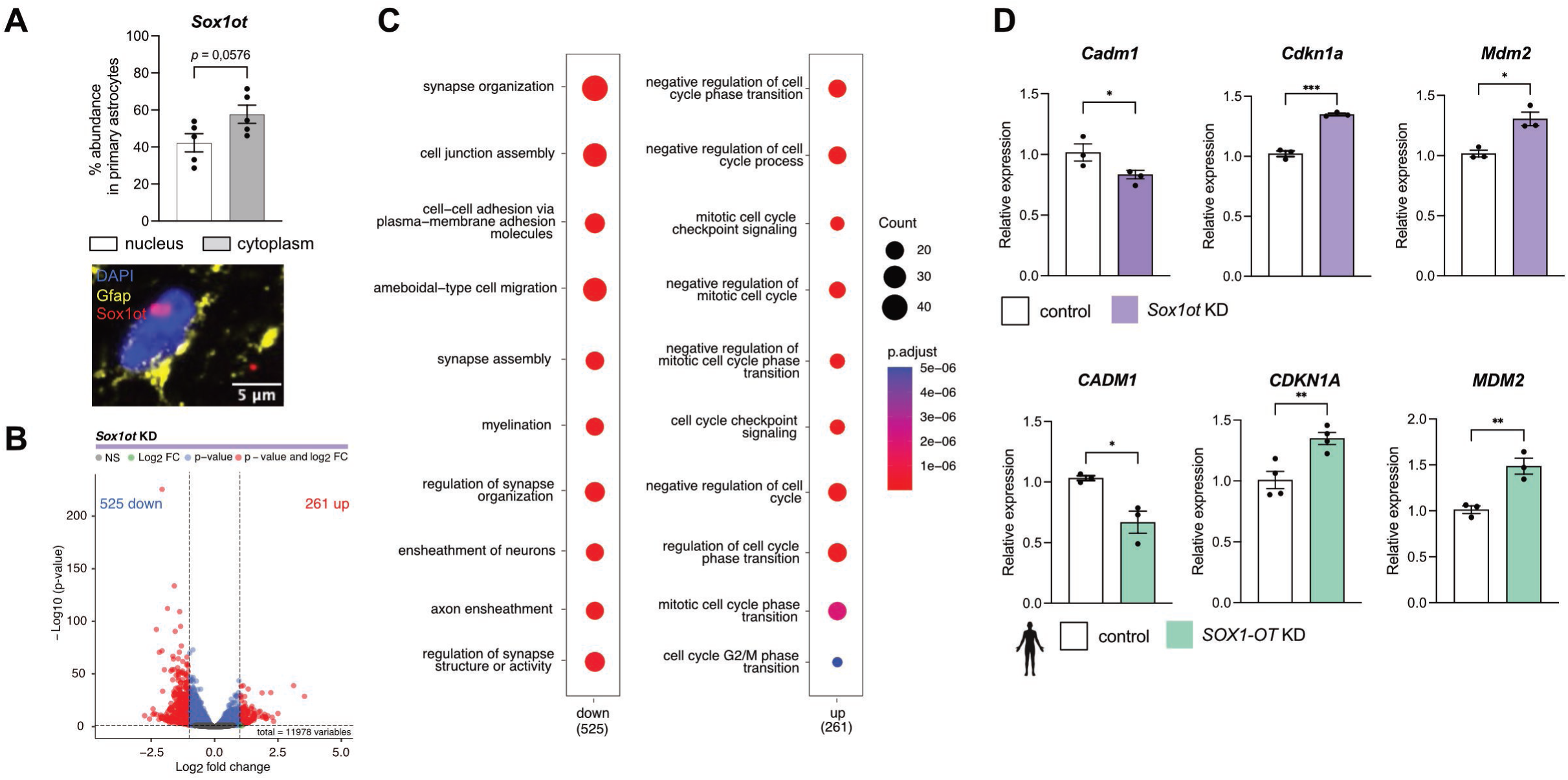
*Sox1ot* knockdown affects synapse organization and cell cycle gene expression in primary astrocytes. **A** *Sox1ot* localisation in astrocytes. Top panel: Bar plot showing qPCR values for *Sox1ot* in nuclear and cytoplasmic fractions prepared from primary astrocytes (unpaired *t-*Test, ****P < 0.0001). Bottom panel: Representative image from the adult mouse cortex showing the RNAscope signal for *Sox1ot* and immunofluorescence for Gfap. Nuclei are stained with DAPI. **B** Volcano plot showing differentially expressed genes from total RNA sequencing of mouse primary astrocytes treated 48 hours with Gapmers to knockdown (KD) *Sox1ot*. Significant genes (adjusted p-value < 0.05 and |log2FC| > 1) are highlighted in red. **C** Plots showing the results of a GO term analysis for selected significant genes displayed in (A). Left panel: Genes down regulated upon *Sox1ot* knockdown. Right panel: Genes up regulated upon *Sox1ot* knock down. Analysis was done using clusterProfiler (v4.6.0). Two-sided hypergeometric test was used to calculate the importance of each term and the Benjamini–Hochberg procedure was applied for the *P* value correction. **D** Top panel: Bar plots showing qPCR results of selected genes for enriched pathways identified in (B) in astrocytes treated 48 hours with Gapmers to knockdown (KD) *Sox1ot* compared to control Gapmers. Bottom panel: qPCR results of selected genes in human pluripotent stem cell-derived astrocytes treated 48 hours with Gapmers to knockdown (KD) human *SOX1-OT* compared to control Gapmers. Gene expression was normalized to 18S. (unpaired *t-*Test; \**P* < 0.05, \*\**P* < 0.01, \*\*\**P* < 0.001). Error bars represent SEM.

To investigate how *Sox1ot* influences gene expression programs underlying the observed functional phenotypes, we performed total RNA sequencing 48 h after *Sox1ot* KD in primary astrocytes. This analysis identified 261 significantly upregulated genes and 525 significantly downregulated genes **(Fig. 3B, Supplemental Table 4)**. Gene Ontology (GO) term analysis showed that the upregulated genes were predominantly associated with negative regulation of the cell cycle, including terms such as “negative regulation of cell cycle transition,” “mitotic cell cycle checkpoint signaling,” and “cell cycle G2/M phase transition,” consistent with the reduced proliferation observed in astrocytes following *Sox1ot* KD **(Fig. 3C, Supplemental Table 5)**. In contrast, the downregulated genes were enriched for GO terms related to synapse organization and cell adhesion, including “synapse organization,” consistent with our findings suggesting impaired synaptic support following *Sox1ot* KD **(Fig. 3C)**. Key genes within these pathways, including *Cell Adhesion Molecule 1* (*Cadm1*; synapse organization) as well as *Cyclin Dependent Kinase Inhibitor 1A (Cdkn1a;* cell cycle regulation) and Mouse Double Minute 2 (*Mdm2*; cell cycle regulation), were validated by qPCR in *Sox1ot*-KD astrocytes **(Fig. 3D)**. Notably, these expression changes were recapitulated in human *SOX1-OT*-KD iPSC-derived astrocytes **(Fig. 3E)**, further supporting a conserved role for *SOX1-OT* in regulating astrocyte gene expression programs across species.

As the gene expression changes observed 48 h after *Sox1ot* KD are likely to reflect a mixture of direct and secondary effects, we used a time-resolved KD approach to identify more direct transcriptional targets and thereby characterize the regulatory role of *Sox1ot* more precisely. To this end, we performed total RNA sequencing following short-term *Sox1ot* KD (8 and 24 h) in primary astrocytes and compared differential gene expression across all time points **(Supplemental Tables 6 and 7**). This analysis identified 39 genes that showed sustained deregulation, suggesting a more direct link to reduced *Sox1ot* levels **(Fig. 4A, Supplemental Table 8)**. GO term analysis showed that the genes with sustained upregulation were enriched for pathways related to cell cycle control, DNA damage response, and particularly p53 signaling, including “mitotic G1/S transition checkpoint signaling,” “DNA damage response,” and “signal transduction by p53 class mediators.” This is notable because p53 is a key regulator of cell cycle progression ^36 12^, which is consistent with the reduced astrocyte proliferation observed following *Sox1ot* knockdown. In contrast, genes with sustained downregulation were enriched for cholesterol and sterol metabolic processes, including “regulation of cholesterol metabolic process” and “sterol metabolic process” **(Fig. 4B, Supplemental Table 9)**.

**Figure 4.**
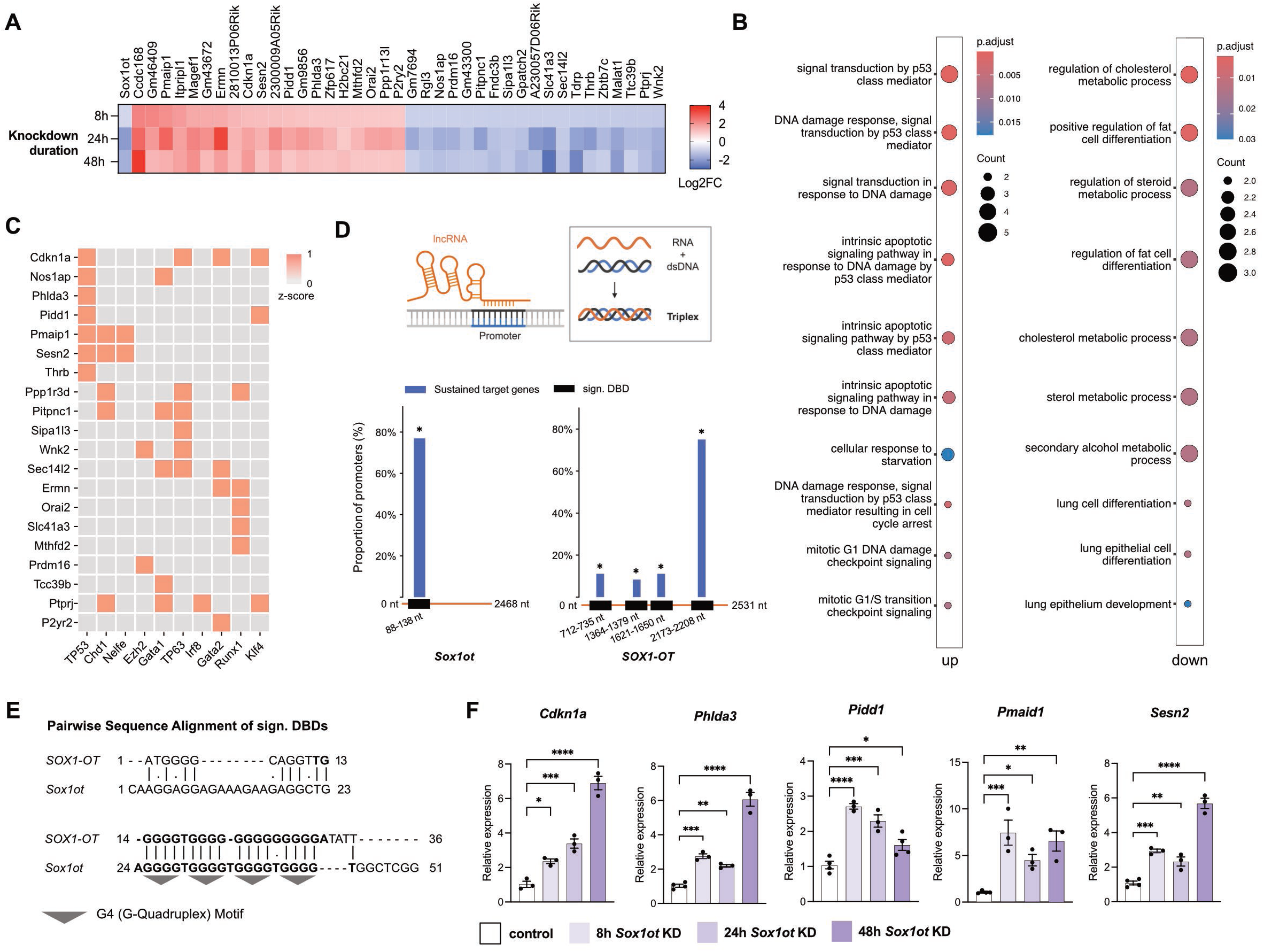
*Sox1ot* target genes identified from a time-resolved knockdown are associated with p53. **A** Heatmap depicting Log2 fold change (FC) of sustained deregulated genes in primary astrocytes treated 8, 24 and 48 hours with Gapmers to knock down (KD) *Sox1ot* identified from total RNA sequencing. **B** Dot plots showing the results of a GO term analysis for genes displayed in (a). Left panel: Genes with sustained up regulation upon 8, 24 and 48 hours *Sox1ot* knock down. Right panel: Genes with sustained down regulation upon 8, 24 and 48 hours *Sox1ot* knock down. Analysis was done using clusterProfiler (v4.6.0). Two-sided hypergeometric test was used to calculate the importance of each term and the Benjamini–Hochberg procedure was applied for the *P* value correction. **C** Clustergram from Enrichr using the ENCODE and ChEA library showing input genes (rows) from (a) and top transcription factors (columns, ranked by combined score). Cell intensity indicates z-score of input gene-transcription factor association. **D** Top panel: Schematic illustration of a triplex-forming domain within a long non-coding (lncRNA, orange) engaging a homopurine stretch in double-stranded DNA (dsDNA, blue-black) via Hoogsteen base pairing in the major groove, forming an RNA–DNA:DNA triple helix (triplex). Bottom panel: Bar charts showing proportion of promoters of sustained deregulated genes identified in (a) with a predicted DNA binding domain (DBD) for *Sox1ot* (left panel) or its human homolog *SOX1-OT* (right panel) from the triplex domain finder tool ^40^. Statistical significance was determined using the Triplex Domain Finder promoter test with Benjamini-Hochberg false discovery rate (FDR) correction of p-values (\**P* < 0.05, FDR-adjusted). LncRNA sequence from 5’ to 3’ in orange, significant DNA-binding domains (sign. DBD) in black with indicated DNA nucleotide (nt) position, sustained target genes in blue. **E** Pairwise sequence alignment of significant DNA binding domains (DBD) identified with the triplex domain finder ^40^ in mouse *Sox1ot* and human *SOX1-OT.* **F** Bar plots showing qPCR results of selected genes identified in (a) with p53 association and predicted *Sox1ot* DNA binding site in primary astrocytes treated 8, 24 and 48 hours with Gapmers to knock down (KD) *Sox1ot*. Gene expression was normalized to 18S (one-way ANOVA; \**P* < 0.05, \*\**P* < 0.01, \*\*\**P* < 0.001, \*\*\*\**P* < 0.0001). Error bars represent SEM.

Next, we performed an Enrichr analysis using the ENCODE and ChEA libraries to identify potential regulators of the genes consistently deregulated upon *Sox1ot* KD. In agreement with the GO term analysis, p53 emerged as the top enriched transcription factor **(Fig. 4C)**, mainly associated with the regulation of genes upregulated following *Sox1ot* KD. Notably, several of these genes are well-established regulators of cell cycle progression, including *Cdkn1a*, a core mediator of cell cycle arrest ^37 38^, which was upregulated upon *Sox1ot* KD. These findings indicate activation of a p53-dependent transcriptional program following *Sox1ot* depletion and suggest a mechanistic link between *Sox1ot* loss, p53 pathway activation, and the observed reduction of astrocyte proliferation. To further test this hypothesis, we took advantage of the fact that nuclear lncRNAs can regulate transcription through direct interactions with DNA, including the formation of RNA-DNA triplex structures ^39^. We therefore investigated whether *Sox1ot* may directly bind to the promoters of genes consistently deregulated upon *Sox1ot* KD. Using the Triplex Domain Finder tool ^40^, we identified a significant DNA-binding domain (DBD) in the 5′ region of mouse *Sox1ot* that was predicted to interact with approximately 80% of the promoters of the persistently deregulated genes. Similarly, a DBD located in the 3′ region of human *SOX1-OT* was predicted to bind nearly 80% of these promoters **(Fig. 4D, Supplemental Table 10)**. Sequence alignment revealed a G-quadruplex motif within the mouse DBD, whereas the human *SOX1-OT* DBD consisted of a G-rich sequence **(Fig. 4E)**.

We next selected five p53 target genes containing predicted *Sox1ot* DNA-binding sites and validated their expression changes upon *Sox1ot* KD by qPCR. Notably, we confirmed increased expression of all selected candidate genes across all knockdown time points **(Fig. 4F)**. Taken together, these findings support a role for nuclear *Sox1ot* in the transcriptional regulation of p53 target genes, likely through direct promoter interactions.

### Effects of *Sox1ot* knockdown on astrocyte cell cycle regulation and p53 activity

Taking into account that p53 is broadly expressed in nearly all nucleated cell types in the human body, these data point to the intriguing hypothesis that *Sox1ot* may act as a regulatory layer that controls the gene-specific activity of p53 in astrocytes. This is consistent with the emerging concept that tissue- and cell type-enriched lncRNAs modulate the activity of ubiquitously expressed transcriptional regulators at context-specific target genes. To further explore this hypothesis, we analyzed key cellular pathways linked to p53 function in response to *Sox1ot* KD, namely the cell cycle, cellular senescence, and apoptosis ^12^.

To assess cell cycle progression in astrocytes following *Sox1ot* KD, we performed propidium iodide DNA staining and flow cytometric analysis of DNA content. The results confirmed our previous observations and revealed a significant arrest of astrocytes in the G1 phase of the cell cycle upon *Sox1ot* knockdown **(Fig. 5A)**. At the same time, cellular senescence, measured by senescence-associated beta-galactosidase (SA-β-gal) activity using the BetaGlo assay **(Fig. 5B)**, and apoptosis, measured using the RealTime-Glo Annexin V Apoptosis and Necrosis assay, remained unchanged **(Fig. 5C)**. These data further support the view that *Sox1ot* regulates specific functions of p53 in astrocytes, particularly astrocyte proliferation.

**Figure 5.**
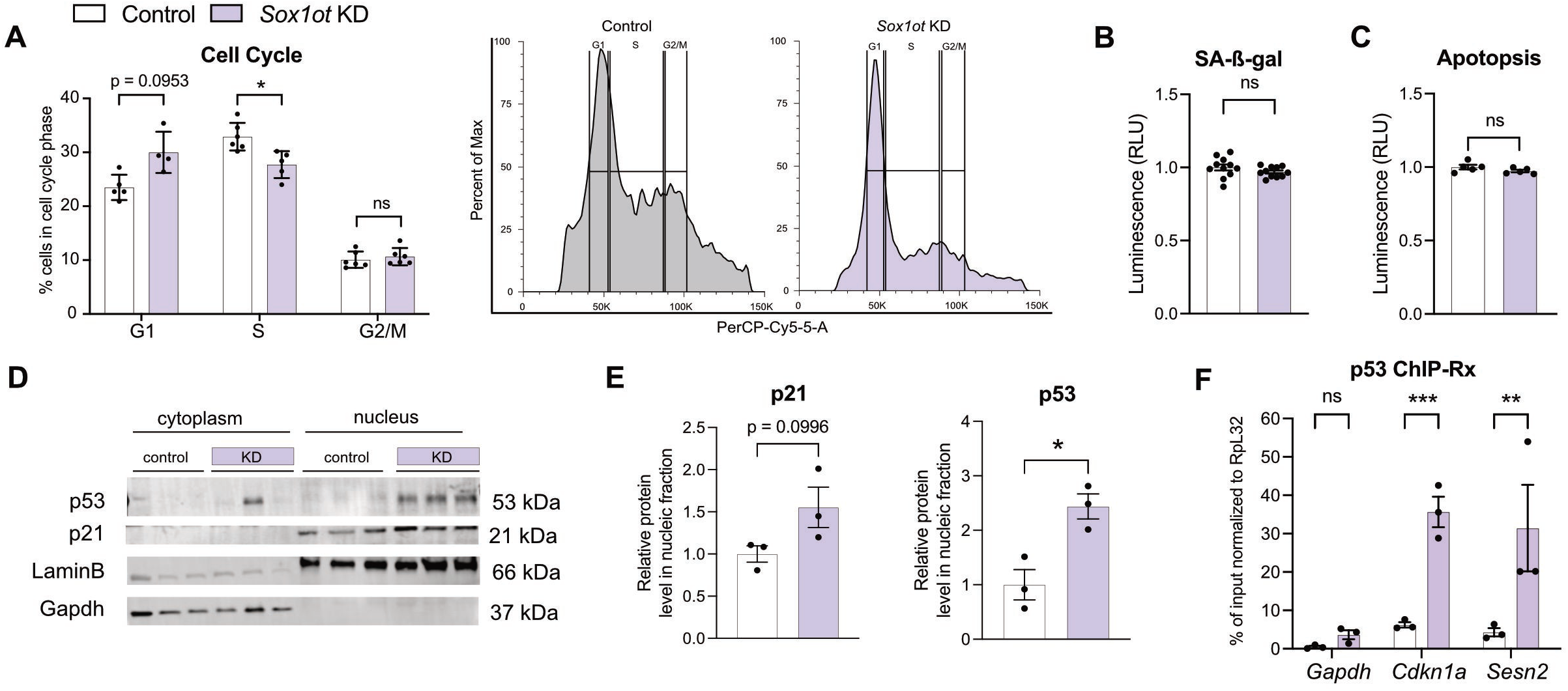
Effects of *Sox1ot* knockdown on astrocyte cell cycle regulation via p53. **A** Left panel: Bar chart depicting the distribution of primary astrocytes across cell cycle phases (G1, S, G2/M) as determined by propidium iodide DNA staining and flow cytometry analysis of DNA content in primary astrocytes treated 24 hours with Gapmers to knock down (KD) *Sox1ot* compared to control Gapmers (one-way ANOVA with Sidak’s multiple comparisons test; \**P* < 0.05). Right panel: Representative flow cytometry histograms depicting the percentage of cells versus PerCP-Cy5 fluorescence intensity (x-axis) for propidium iodide used in the cell cycle assay in (a). **B** Bar plot showing results of senescence assay quantifying senescence-associated β-galactosidase (SA-β-gal) levels in relative light units (RLU) in primary astrocytes treated 24 hours with Gapmers to knock down (KD) *Sox1ot* compared to control Gapmers (unpaired *t-*Test; ns not significant). **C** Bar plot showing results of apoptosis assay quantifying Annexin V levels in relative light units (RLU) in primary astrocytes treated 24 hours with Gapmers to knock down (KD) *Sox1ot* compared to control Gapmers (unpaired *t-*Test; ns not significant). **D** Representative Western blots of p53, p21 and loading controls LaminB and Gapdh in nuclear fractions of primary astrocytes treated 24 hours with Gapmers to knockdown (KD) *Sox1ot* compared to control Gapmers (control). Molecular weights (kDa) are indicated. **E** Bar plots depicting densitometric quantification of p21 or p53 protein levels normalized to LaminB from (D), (unpaired *t-*Test; \**P* < 0.05). **F** Bar plot showing qPCR results of p53 occupancy at target gene promoter sites following ChIP-Rx in primary astrocytes treated 24 hours with Gapmers to knockdown (KD) *Sox1ot* compared to control Gapmers. Data normalized to drosophila RpL32 (two-way ANOVA with Sidak’s multiple comparisons test; \*\**P* < 0.01, \*\*\**P* < 0.001, ns not significant). Error bars represent SEM.

A major p53 target mediating G1 cell cycle arrest is the *Cdkn1a* gene, which encodes the p21 protein ^38^. Since we had already demonstrated that loss of *SOX1-OT/Sox1*ot leads to increased *Cdkn1a* levels, we next quantified p53 and p21 protein levels following *Sox1ot* knockdown by western blot. In line with our previous findings, we observed significant increases in nuclear p21 and p53 levels in astrocytes following *Sox1ot* KD **(Fig. 5D,E)**.

To directly test whether *Sox1ot* modulates p53 binding to target gene promoters involved in cell cycle regulation in astrocytes, we performed quantitative p53 chromatin immunoprecipitation with exogenous spike-in normalization (ChIP-Rx). This approach allowed us to account for changes in p53 protein levels during *Sox1ot* knockdown by using spike-in chromatin and antibodies derived from *Drosophila melanogaster*. Following *Sox1ot* KD, we observed a significant increase in p53 occupancy at the promoter regions of key target genes, including *Cdkn1a* and *Sesn2* (≈20–50% over input) **(Fig. 5F)**. *Sesn2* is another p53 target gene implicated in the regulation of cell cycle arrest through inhibition of mTORC1 signaling and suppression of STAT3 activity ^41^. IN lien with these findings, we observed that CRISPRa-mediated overexpression of *Sox1ot* in astrocytes increased proliferation and decreased the number of cells in the G1 phase **(Fig. S4)**.

Together, these data support a model in which *Sox1ot* restrains p53 occupancy at cell-cycle regulatory genes, thereby limiting p53-dependent G1 arrest and maintaining astrocyte proliferative capacity.

## Discussion

In this study, we identify *SOX1-OT/Sox1ot* as a conserved, brain-enriched lncRNA that is downregulated in AD and in reactive astrocytes *in vitro*, and show that it regulates key astrocyte support functions while restraining a p53-dependent cell-cycle arrest program. Notably, the only previous mechanistic study of *SOX1-OT* in the brain described a distinct role in human neuronal differentiation, in which *SOX1-OT* promoted neurogenesis ^29^, suggesting that *SOX1-OT* may exert context-dependent functions in the developing and adult brain.

Astrocytes must dynamically balance proliferative competence, inflammatory responsiveness, metabolic support, and synaptic regulation making them particularly dependent on regulatory mechanisms that tailor broadly expressed pathways to cell-type-specific needs ^4^. *Sox1ot* KD in mouse astrocytes and *SOX1-OT* KD in human iPSC-derived astrocytes consistently impaired glutamate uptake, lactate secretion, and astrocyte support functions. However, these functional deficits may in part reflect secondary consequences of the transcriptional state change induced by *SOX1-OT* loss, rather than immediate primary targets of the lncRNA. In this regard, our time-resolved analysis is particularly informative, as it identified a smaller set of sustained early transcriptional changes that converged strongly on p53 targets and cell-cycle inhibitory pathways. In fact, computational analysis predicted triplex-forming DNA-binding domains in both mouse *Sox1ot* and human *SOX1-OT* that could interact with the promoters of approximately 80% of these persistently deregulated genes. This interpretation is conceptually consistent with temporally resolved lncRNA perturbation studies showing that early measurements help distinguish primary regulatory events from later secondary transcriptional cascades ^42^.

Functionally, we observed increased expression of p53 target genes, including *Cdkn1a*, together with elevated levels of its protein product p21. As p21 inhibits cyclin-CDK complexes required for G1/S progression ^12^, these changes provide a plausible mechanistic link between enhanced p53 activity and the G1 arrest observed after *Sox1ot* depletion. In line with this model, increased p53 occupancy at the *Cdkn1a* promoter was accompanied by a shift toward G1 arrest.

This framework is particularly relevant because p53 is a pleiotropic stress-response factor that can induce cell-cycle arrest, senescence, apoptosis, and metabolic adaptation depending on cellular context ^10 11 12^. How these different outputs are selected in specific cell types remains poorly understood. Our findings suggest that *SOX1-OT* contributes to this specificity in astrocytes and are consistent with the emerging view that lncRNAs fine-tune context-dependent gene regulatory programs rather than acting as simple on/off switches ^8 43 44 9 45^. Together with the increased p53 promoter occupancy observed after *Sox1ot* depletion, these findings support a model in which *SOX1-OT* normally restrains access or recruitment of p53 to a subset of target loci in astrocytes **(Fig. 6)**.

**Figure 6.**
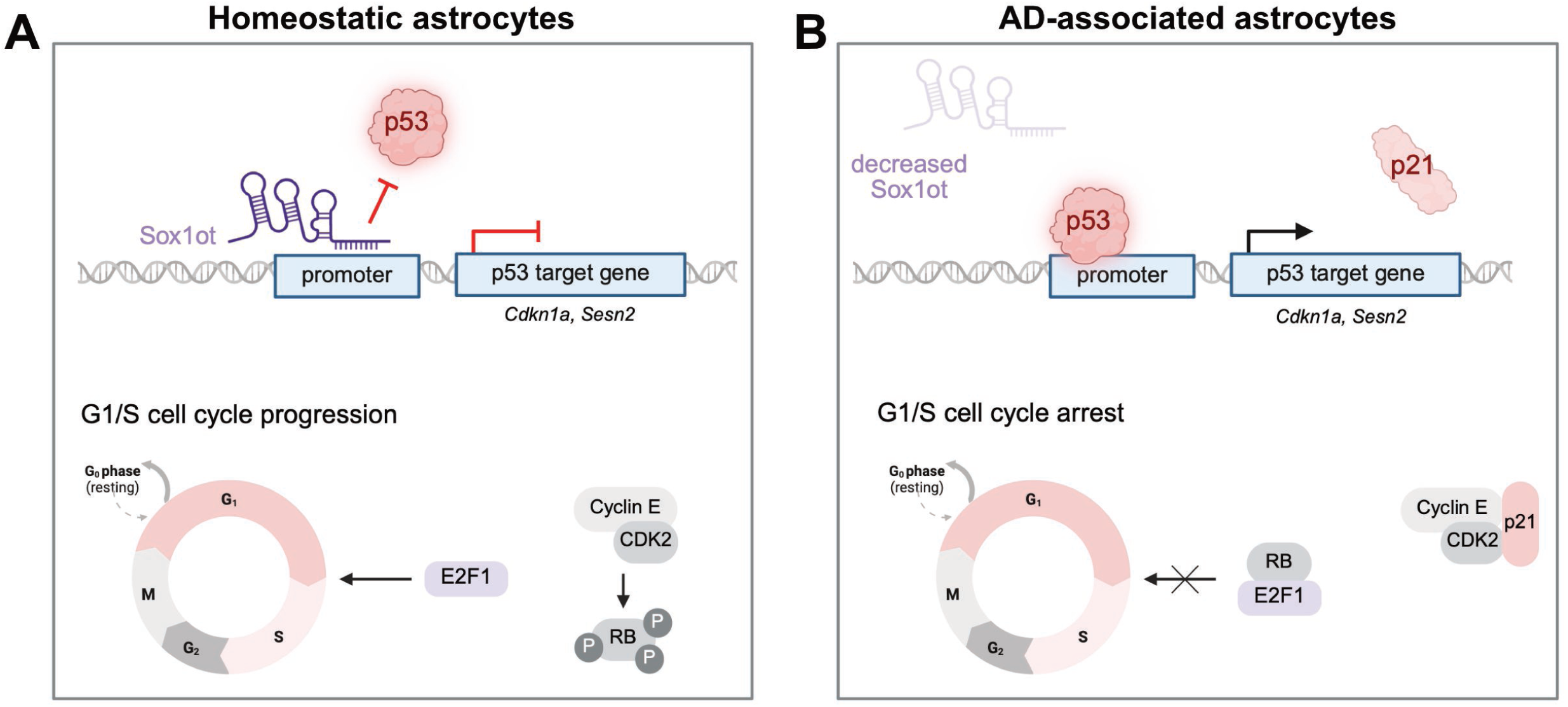
*Sox1ot* mechanism of action in astrocytes. **A** Schematic illustration of the *Sox1ot* mechanism of action in the nucleus of homeostatic astrocytes. *Sox1ot* binds to the promoter region of p53-target genes involved in cell cycle regulation, keeping p53 away and thereby inhibiting gene expression while ensuring G1/S cell cycle progression and maintaining proliferative capacity. E2F transcription factor 1 (E2F1) facilitates the progression from G1 into S phase by regulating the expression of genes essential for S-phase entry, DNA synthesis, and mitosis. E2F1 activity is tightly regulated by retinoblastoma protein (RB), which binds E2F1 to inhibit it until phosphorylation (P) by cyclin-dependent kinases (CDKs) releases it. **B** Schematic illustration of the effect of *Sox1ot* loss in the nucleus of astrocytes observed in Alzheimer’s disease (AD) or reactive astrocytes. *Sox1ot* loss at the promoter sites of p53-target genes involved in cell cycle regulation increases p53 promoter occupancy and leads to increased expression of e.g. Cyclin-dependent kinase inhibitor 1A (*Cdkn1a*) mRNA and subsequent p21 protein levels. P21 binds to the complex of Cyclin E and CDK2, which impairs phosphorylation of RB. RB in turn binds to and inhibits E2F1 activity, resulting in G1/S cell cycle arrest.

This is consistent with the biology of astrocytes, which, unlike post-mitotic neurons, retain proliferative capacity and can re-enter the cell cycle in response to CNS injury, where they contribute to reactive remodeling and glial scar formation ^4^. By contrast, we did not observe corresponding changes in apoptosis or SA-β-gal activity, indicating that under the employed experimental conditions *SOX1-OT* loss does not simply trigger a full canonical p53 program, but rather promotes a more selective p53-dependent transcriptional response centered on cell-cycle control.

The disease relevance of this mechanism is supported by several converging observations. *SOX1-OT* was reduced in human AD brain tissue, in public AD datasets, and both mouse *Sox1ot* and human *SOX1-OT* were downregulated in cytokine-induced reactive astrocyte states. Astrocyte reactivity is increasingly recognized as a central feature of AD and other neurodegenerative diseases ^3 4^, where astrocytes respond to proteinopathy but can also become active contributors to neuroinflammation and synaptic dysfunction. To model this transition *in vitro*, we used the IL-1α/TNF-α/C1q cytokine cocktail originally described by Liddelow and colleagues, which has been widely adopted to induce a neurotoxic reactive astrocyte state ^17^. Although this paradigm does not recapitulate the full complexity of AD, it captures a widely used inflammatory astrocyte state with disease relevance. Against this background, the consistent downregulation of *SOX1-OT/Sox1ot* in both mouse and human astrocytes supports the idea that loss of this lncRNA may be part of a broader astrocyte state transition associated with neuroinflammatory or neurodegenerative conditions.

Importantly, *Sox1ot* depletion reproduced several features associated with dysfunctional astrocyte states, including impaired glutamate handling, reduced metabolic support, inflammatory gene induction, and activation of a p53-linked cell-cycle arrest program. However, our data do not support the conclusion that *Sox1ot* loss alone is sufficient to induce a fully established senescence phenotype. Instead, the observed changes are more consistent with a selective, stress-associated, pre-senescent state in which p53-dependent cell-cycle control is engaged without overt apoptosis or robust SA-β-gal induction. This interpretation is in line with growing evidence that astrocyte dysfunction in AD spans a spectrum of partially overlapping reactive and senescence-like states ^46 4^.

More broadly, these findings support the idea that astrocyte-enriched lncRNAs can act as molecular filters that shape the target selectivity of widely expressed regulatory pathways. In this regard, *SOX1-OT* may function less as a general on/off regulator of p53 and more as a molecular filter that biases p53 activity toward or away from specific transcriptional programs. This view fits with the broader emerging principle that lncRNAs contribute to cellular identity not only through lineage-restricted expression, but also by shaping the target selectivity of widely expressed transcriptional and chromatin regulators ^47 8 43 44 9 45 6^.

Several limitations of the present study should be considered. Although we analyzed the role of *SOX1-OT/Sox1ot* in primary mouse astrocytes and human iPSC-derived astrocytes, several studies have pointed to substantial heterogeneity among astrocyte subtypes across brain regions and disease stages ^46 4^. A more detailed analysis of *SOX1-OT* expression in the human brain using snRNA-seq methods that robustly capture lncRNA expression ^8^, or complementary histological approaches, will therefore be important to define its expression pattern in the healthy and diseased brain. This would also help address a current technical limitation of our study, namely that direct assessment of *SOX1-OT* dysregulation in astrocytes from AD patient single-nucleus datasets. This remains difficult, likely because lncRNAs such as *SOX1-OT* are incompletely captured in currently available 3′-biased snRNA-seq datasets ^34^. Moreover, while our computational and ChIP-based analyses support a promoter-associated nuclear function of *SOX1-OT*, the proposed RNA-DNA interaction mechanism remains inferential and should be tested directly in future work. Finally, although we focused here on astrocyte phenotypes, our data indicate that *SOX1-OT* is also expressed in other neural cell types in the human brain, particularly OPCs. It will therefore be important to determine whether *SOX1-OT* has additional functions outside astrocytes.

In conclusion, our study identifies *SOX1-OT* as a conserved regulator of astrocyte biology with a mechanistic link to p53-dependent cell-cycle control. We propose that *SOX1-OT* provides a lncRNA-based mechanism for adapting the pleiotropic p53 pathway to the specific functional requirements of astrocytes. Its downregulation in AD and in reactive astrocytes suggests that loss of this regulatory layer may contribute to maladaptive astrocyte state transitions in neurodegeneration. Future studies should determine whether restoring *SOX1-OT* expression, or targeting the downstream p53 program it constrains, can normalize astrocyte function in human disease-relevant models. Given the increasing feasibility of RNA-targeting therapeutic strategies, including antisense oligonucleotides and RNA-based modulation approaches, *SOX1-OT* and its downstream p53 regulatory axis may represent a tractable entry point for modulating astrocyte dysfunction in neurodegenerative disease.

## Acknowledgements

AF was supported by the DFG (Deutsche Forschungsgemeinschaft) priority program 2502 EPIADAPT, SFB1286. The German Federal Ministry of Research, Technology and Space (BMFTR) via the ERA-NET Neuron project EPINEURODEVO; The EU Joint Programme- Neurodegenerative Diseases (JPND)-EPI-3E; Germany’s Excellence Strategy-EXC 2067/1 390729940. FS was supported by the GoBIO project miRassay (16LW0055) by the BMFTR. ID was supported by NIH RF1AG078299 & RO1 AG084624. UF was supported by the Hans und Ilse Breuer Stiftung and the International Max-Planck Research School (IMPRS) Genome Science Göttingen.

## Availability of Data and Materials

RNA-sequencing data is available via the GEO database: GSE330559 https://www.ncbi.nlm.nih.gov/geo/query/acc.cgi?acc=GSE330559

Enter token **cxmvkugcrterlsr**

## Author contributions

UF planned and conducted the majority of the experiments. TP and DMK helped with statistical analysis. NH helped with the analysis of tissue and brain-specific lncRNAs, VG and TT helped with experiments. SS, SB and ALS coordinated and performed RNAseq experiments. MG, TP, JC, FS performed analysis of APP mice. AF conceived the project, supervised progress. UF and AF wrote the manuscript.

