## Supplemental figures and tables for "Loss of the lncRNA *SOX1-OT* promotes p53-dependent cell-cycle arrest in astrocytes": Supplementary Figures 1-4.docx

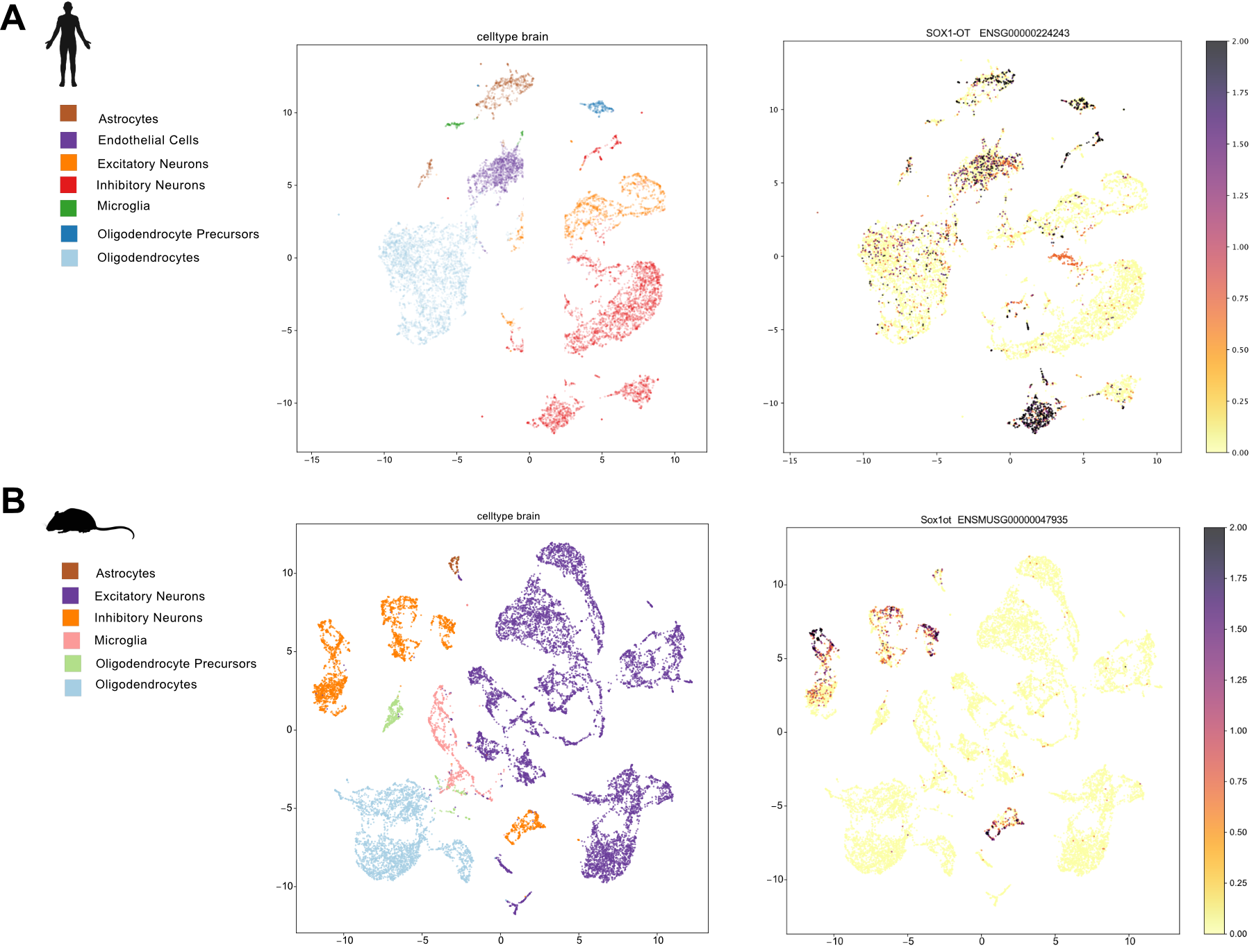

**Figure S1**

***SOX1-OT* and *Sox1ot* detection in snRNAseq data of the healthy human or mouse brain.**

**A** *SOX1-OT* expression in the human brain. Data was obtained from snRNAseq data (iCELL8 technology) from healthy human brains (prefrontal cortex, BA9) {Schröder, 2024}. Right panel: UMAP clustering based on human snRNAseq data. Left panel: UMAP clustering as depicted in the right panel showing the expression of *SOX1-OT*. **B** *Sox1ot* expression in the mouse brain. Data was obtained from snRNAseq data (10X technology) from healthy mice (hippocampus) {Michurina, 2021}. Right panel: UMAP clustering based on mouse snRNAseq data. Left panel: UMAP clustering as depicted in the right panel showing the expression of *Sox1ot*.

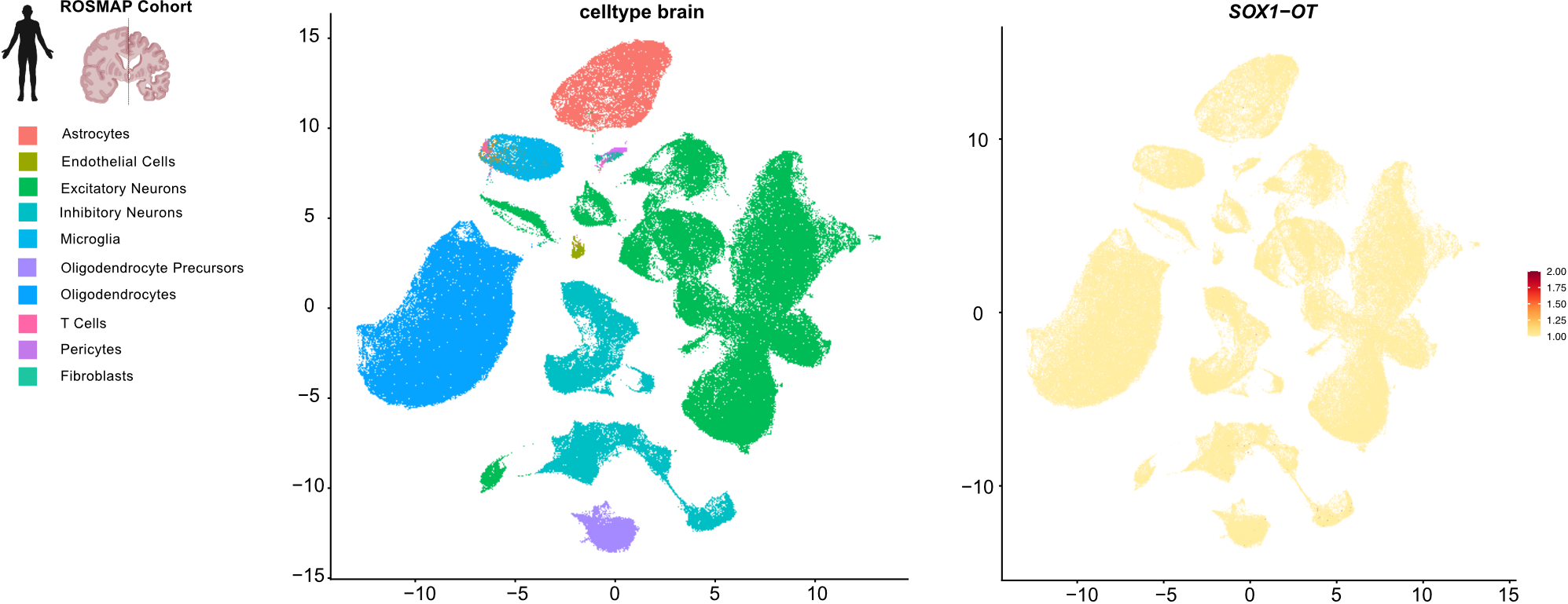

**Figure S2**

***SOX1-OT* detection in snRNAseq data from the ROSMAP cohort.**

*SOX1-OT* expression in the human brain. Data was obtained from snRNAseq data (10X Genomics) from healthy human brains and Alzheimer’s disease patient brains from the ROSMAP cohort {Mathys, 2023}. Right panel: UMAP clustering based on human snRNAseq data. Left panel: UMAP clustering as depicted in the right panel showing the expression of *SOX1-OT*.

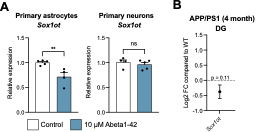

**Figure S3**

***Sox1ot* expression changes in models of Alzheimer’s disease.**

**A** Bar plots showing *Sox1ot* expression upon 24 hour treatment with 10 µM amyloid beta (Abeta)1-42 protofibrils compared to DMSO vehicle control (control) within the insert co-culture system in primary astrocytes (left) or primary cortical neurons (right). Gene expression was normalised to 18S (unpaired *t* test, ***P* < 0.01, ns = not significant). Error Bars represent SEM. **B.** Dot plots showing Log2 fold change (FC) from total RNA sequencing data of *Sox1ot* expression in the dentate gyrus (DG) of 4 month old APP/PS1 mice compared to wild type (WT) controls, n = 4 animals per group (Wald Test,*P < 0.1*).

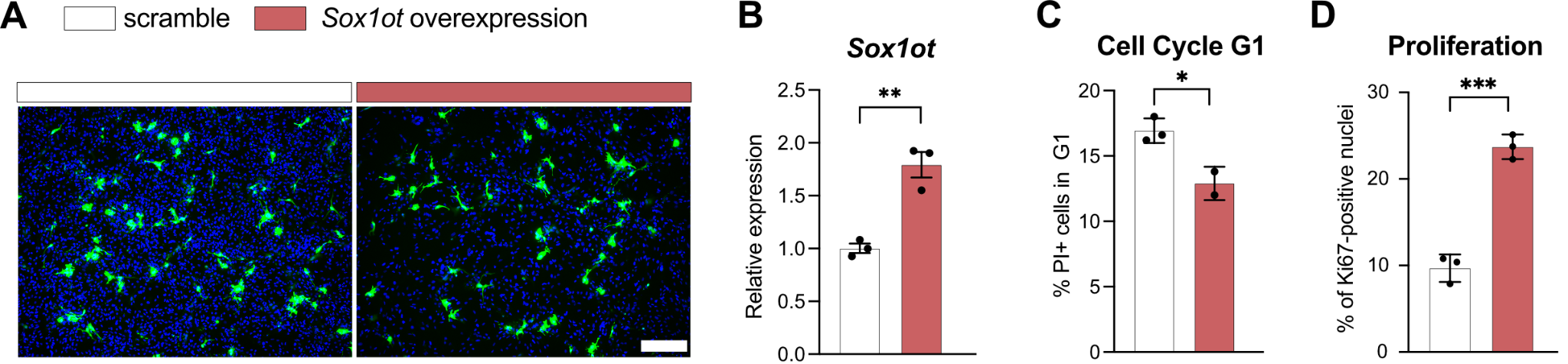

**Figure S4**

**CRISPRa-mediated overexpression of *Sox1ot* in primary astrocytes.**

**A** Representative images of 72 hour CRISPRa-mediated overexpression of *Sox1ot* compared to scramble control guide RNA in primary astrocytes. Successfully transfected cells with GFP-signal in green. Nuclei were stained with DAPI in blue. **B** Bar plots showing *Sox1ot* levels after 72 hour CRISPRa-mediated *Sox1ot* overexpression compared to scramble control. Gene expression was normalized to 18S. **C.** Bar chart depicting the distribution of primary astrocytes across cell cycle phase G1 as determined by propidium iodide DNA staining and flow cytometry analysis of DNA content in primary astrocytes treated 74 hours with CRISPRa guide RNA to overexpress *Sox1ot* compared to scramble control guide RNA (unpaired *t-*Test; **P* < 0.05). **D.** Bar chart showing the relative amount of proliferating primary astrocytes determined by Ki67 immunofluorescence imaging 72 hours after CRISPRa-mediated *Sox1ot* overexpression in comparison to scramble control guide RNA (unpaired *t-*Test; ***P* < 0.01). Error bars represent SEM.
